# Macrophages protect against sensory axon degeneration in diabetic neuropathy

**DOI:** 10.1101/2024.01.30.577801

**Authors:** Sara Hakim, Aakanksha Jain, Veselina Petrova, Jonathan Indajang, Riki Kawaguchi, Qing Wang, Elif Sude Duran, Drew Nelson, Stuart S. Adamson, Caitlin Greene, Clifford J. Woolf

## Abstract

Diabetic peripheral neuropathy (DPN) is a common complication of diabetes, causing sensory loss and debilitating neuropathic pain^1,2^. Although the onset and progression of DPN have been linked with dyslipidemia and hyperglycemia^3^, the contribution of inflammation in the pathogenesis of DPN has not been investigated. Here, we use a High Fat High Fructose Diet (HFHFD) to model DPN and the diabetic metabolic syndrome in mice. Diabetic mice develop persistent heat hypoalgesia after three months, but a reduction in epidermal skin innervation only manifests at 6 months. Using single-cell sequencing, we find that CCR2+ macrophages infiltrate the sciatic nerves of diabetic mice well before axonal degeneration is detectable. We show that these infiltrating macrophages share gene expression similarities with nerve crush-induced macrophages^4^ and express neurodegeneration-associated microglia marker genes^5^ although there is no axon loss or demyelination. Inhibiting this macrophage recruitment in diabetic mice by genetically or pharmacologically blocking CCR2 signaling results in a more severe heat hypoalgesia and accelerated skin denervation. These findings reveal a novel neuroprotective recruitment of macrophages into peripheral nerves of diabetic mice that delays the onset of terminal axonal degeneration, thereby reducing sensory loss. Potentiating and sustaining this early neuroprotective immune response in patients represents, therefore, a potential means to reduce or prevent DPN.

## Main Text

Peripheral neuropathy is a debilitating neurodegenerative condition where the dying back of distal axonal endings of those long sensory axons that innervate the skin of the hands and feet produces a “glove and stocking” pattern of sensory loss^1,3^. 537 million people worldwide had diabetes in 2021 and approximately 50% of these patients will eventually develop diabetic peripheral neuropathy (DPN)^6,7^. About 30-40% of those patients with DPN also have neuropathic pain, which typically manifests as spontaneous burning pain in the denervated locations^1,3,7,8^.

Axonal degeneration in the peripheral nervous system has been most extensively studied in the context of traumatic nerve injury. Following a nerve transection or crush injury, the distal axons of PNS neurons undergo Wallerian degeneration^9,10^. Nerve injury also leads to the recruitment of immune cells both directly at the injury site and in the distal portion of the axons^4,11^.

What is currently unclear is whether immune cells contribute either to axonal degeneration or disease progression in peripheral neuropathy. Understanding if and how the immune system may contribute to diabetic peripheral neuropathy will allow for the potential development of novel therapies targeting the neuro-immune axis for the treatment of diabetic neuropathy.

To study the neuro-immune mechanisms of diabetic neuropathy, we have modeled type 2 diabetes and its associated metabolic syndrome in mice. The incidence of type 2 diabetes is increasing as a direct result of the heightened prevalence of obesity and the consumption of high-caloric food (Western diet)^12,13^. To mimic the consumption of high-caloric food, we fed wild-type (WT) male mice on a high fat diet (60% Kcal from lard) supplemented with high fructose water (42g/L D-(-)-Fructose) (HFHFD) (Fig. 1a). This diet leads both to a consistent weight gain (Fig.1b) and to glucose intolerance (Fig.1c) when compared to a low-fat control diet supplemented with plain water (CD).

**Figure 1.**
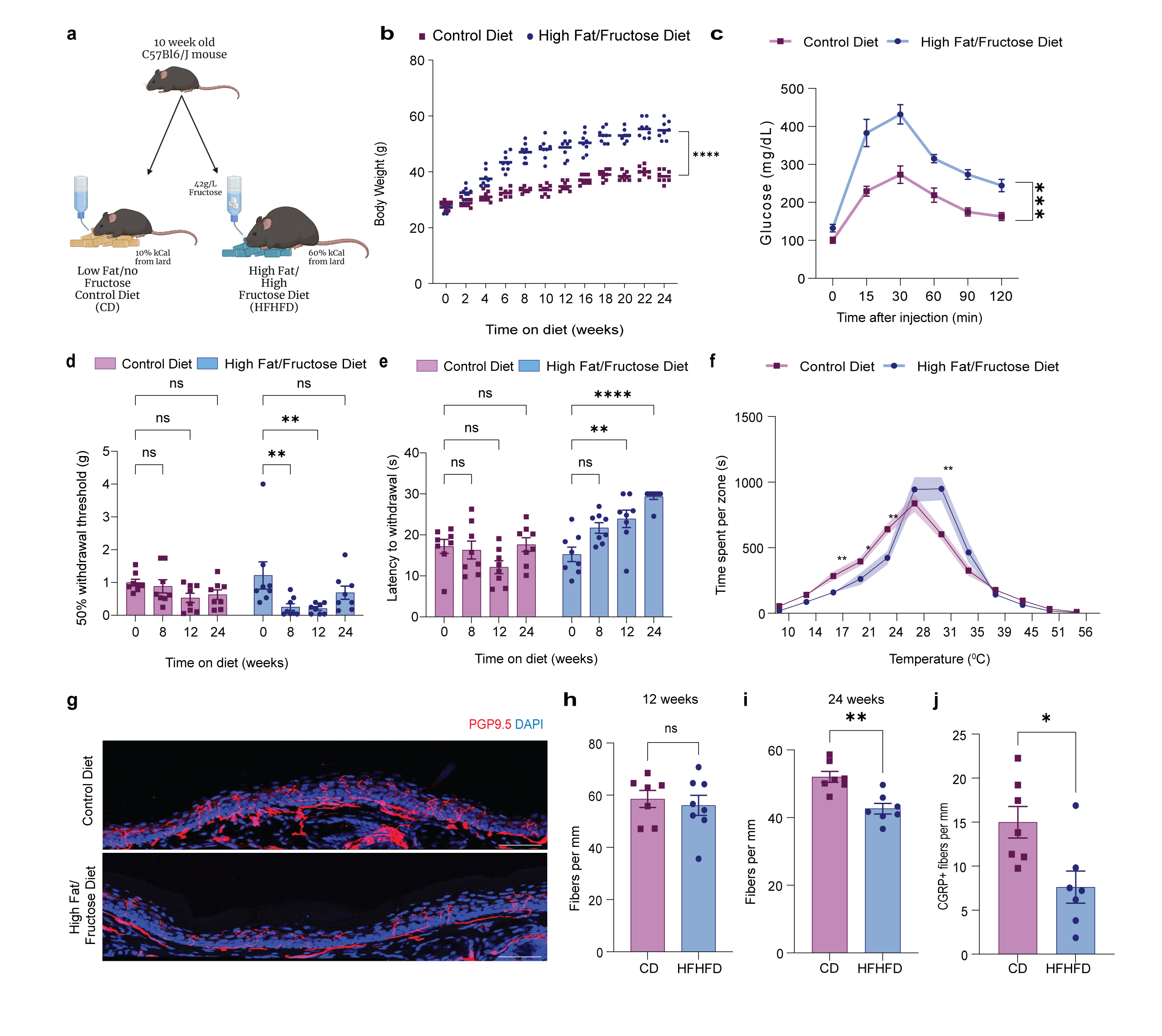
A High Fat High Fructose diet leads to obesity, diabetes, and peripheral neuropathy. **a.** Illustration of the high fat high fructose diet (HFHFD) vs control diet (CD) administered to induce diabetes. **b.** Body weights of male C57Bl6 mice fed HFHFD and CD over six months. **c.** Glucose tolerance tests after fasting on mice fed HFHFD or CD (p-value <0.0001). **d.** Mechanical withdrawal thresholds of mice fed HFHFD or CD over time using an up down von Frey method (p-value HFHFD 0 vs 8w =0.0052, and 0 vs 12w = 0.0032). **e.** Heat withdrawal latency of mice fed HFHFD or CD over time using the Hargreaves assay (p-value HFHFD 0 vs 12w = 0.0068 and 0 vs 24w<0.0001). **f.** Time spent on a thermal gradient ring of mice fed HFHFD or CD for 12 weeks (n=8 mice per group) (p-value at 17C = 0.007791, at 21C = 0.036572, at 24C= 0.007500, at 31C = 0.001774). **g.** Representative PGP9.5 staining of hind paw skin of mice fed HFHFD or CD for 24 weeks showing nerve fibers in the epidermis marked by PGP9.5 and cell layers by DAPI+. **h.** Quantification of intraepidermal nerve fiber density at 12 weeks of diet, and **i.** 24 weeks of diet (p-value =0.0013). **j.** and quantification of CGRP+ fibers at 24 weeks of diet**(p-value =0.0139).**

### High Fat High Fructose Diet leads to diabetic neuropathy

Diabetic mice develop a pain-like behavior in response to innocuous mechanical stimuli (allodynia) starting at 8 weeks (Fig. 1d) and a reduction in responsiveness to noxious heat (heat hypoalgesia), first detectable at 12 weeks (Fig. 1e) which is a common feature in patients with diabetic neuropathy^14^, and in mice^15,16^. We evaluated whether there are sex differences in the response to this HFHFD and found that female mice also gain weight significantly over time on the HFHFD compared to the CD (Fig. S1a) and develop significant mechanical allodynia and heat hypoalgesia similar to male mice (Fig. S1b,c). We then continued the characterization with male mice. To further assess changes in thermal sensitivity we used an unsupervised thermal gradient ring assay to evaluate thermal preferences and found that diabetic mice have a higher tolerance to hot regions of the ring, and more cool avoidance (Fig. 1f, S2a). The mechanical allodynia, unlike the heat hypoalgesia, fully resolved by 24 weeks on the diet, suggesting that the dysfunction driving this phenomenon disappears as the disease progresses (Fig. 1d).

Reduction in skin intraepidermal nerve fiber density (IENFD) is the driver of DPN diagnosis in patients^7^. Mice fed with the HFHFD do not develop reduced skin IENFD at 12 weeks (Fig. 1h) or demyelination in the nerves (Fig. S2), but terminal axon degeneration is clear after 24 weeks on the diet (Fig. 1g,1i). More specifically, we found that CGRP+ fibers in the epidermis of the skin degenerate in these mice at 24 weeks, indicating that this is a small fiber neuropathy (Fig. S2b,1j), exactly as seen in patients.

We next asked whether the delayed terminal axon degeneration in these mice is Sarm1-dependent, as is Wallerian degeneration^9,10,17–19^. We fed Sarm1-KO mice on the HFHFD and found that absence of Sarm1 prevents a loss of intraepidermal fibers in the skin at 24 weeks (Fig. S3a). Interestingly, the Sarm1-KO mice also did not develop the heat hypoalgesia seen in WT mice at 12 weeks on a HFHFD (Fig. S3b). This suggests that even though there is no detectable reduction of IENFD at this stage, the loss of noxious heat sensitivity reflects SARM1-dependent axonal dysfunction. Sarm1-KO mice did, however, develop mechanical allodynia at 8 weeks on the HFHF diet, much like WT mice. Additionally, we found that in Sarm1-KO mice the mechanical allodynia did not resolve at 24 weeks on the diet as it does in WT mice (Fig. S3c) suggesting that the recovery from the mechanical allodynia in the WT mice at 24 weeks may be a consequence of a late mechanosensitive axon terminal degeneration. These data show that a Western diet comprised of high fat and high fructose leads to obesity, diabetes, and sensory neuropathy, capturing multiple features of the clinical disease. We also find that behavioral changes are detectable well before axon degeneration, and that the heat sensitivity loss as well as the axonal degeneration in the skin are both Sarm1-dependent, whereas mechanical allodynia is not.

### Macrophages are recruited into nerves of diabetic mice

Diabetes and obesity are accompanied by a metabolic syndrome, where the immune system is implicated in disease pathogenesis^20^. To evaluate if neuro-immune interactions contribute to DPN, we sought to identify changes in immune cells that may directly affect sensory neurons at 12 weeks of feeding on the HFHFD when mice develop heat hypoalgesia but before axon terminal degeneration is observed. Immune cells reside in proximity to sensory neurons both in nerves and the skin^4,21^. To determine whether there are changes in immune cells in close proximity to sensory neurons after 12 weeks on the diet, we FACS-sorted CD45+ immune cells from the hind paw skin and sciatic nerve and used single cell sequencing to evaluate immune cell composition (Fig. 2a, S7a), analyzing 8462 cells in the sciatic nerve (Fig. S4a) and 9793 cells in the skin (Fig. S7b).

**Figure 2.**
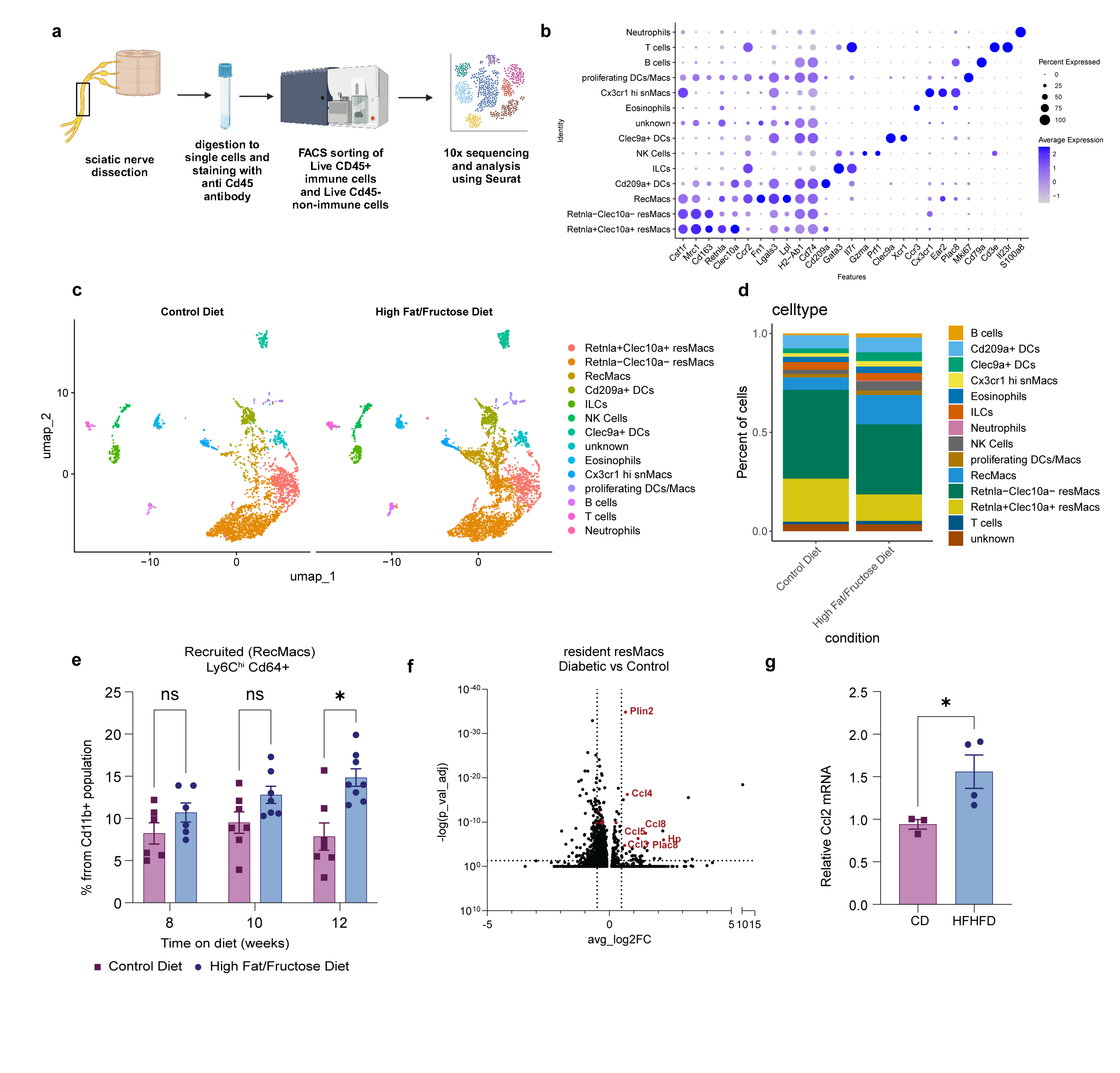
CCR2+ macrophages infiltrate sciatic nerves of diabetic mice and resident macrophages become pro-inflammatory. **a.** Workflow for tissue collection, processing, sorting, and sequencing of sciatic nerve immune cells from mice fed HFHFD or CD for 12 weeks. **b.** Dot plot of marker genes used to identify different immune cells. **c.** UMAP plot showing clusters identified in sciatic nerves of CD and HFHFD fed mice (2 samples per group). **d.** Bar plot of cell proportions showing increase in CCR2+ macrophages in sciatic nerves of HFHFD vs CD-fed mice. **e.** Flow cytometry quantification of nerves of mice fed CD or HFHFD for different durations assessing recruited macrophages (Ly6C^hi^ Cd64+) in Cd11b+Cd45+ live cell gate (p-value at 12w = 0.0123). **f.** Volcano plot of gene markers that differentially expressed in resident (Csf1r+Ccr2-) sciatic nerve macrophages (resMacs) clusters in HFHFD-fed mice compared to CD-fed mice showing increased chemokines and cytokines. **g.** RT-qPCR quantification of Ccl2 in nerves from mice fed HFHFD and CD for 12 weeks (p-value = 0.0469).

The sciatic nerve immune dataset enabled an in-depth analysis of changes in the nerve immune landscape in diabetic neuropathy, which has not been examined before. Recently, another group performed a study in high-fat diet-fed mice^22^, but did not enrich for immune cells, and in consequence did not capture the depth and heterogeneity of our dataset. We identified 13 clusters of immune cells in the sciatic nerve, including five distinct sets of macrophages defined by expression of *Csf1r*, including 2 sciatic nerve resident macrophages (resMacs) which were *Ccr2-*, of which one was *Retnla+Clec10a*+ and the other *Retnla-Clec10a-* and are likely the epineural and endoneurial populations previously described^4^. We also identified a small cluster of resident macrophages *which were Cx3cr1* high, similar to macrophages in the skin that are thought to play a role in axon growth^21^. In addition, we identified two dendritic cell clusters (DCs) that express high MHCII (*H2-Ab1*) and *Cd74*, one of which expresses *Cd209a* and the other, much likecDC1s, with expression of *Clec9a* and *Xcr1*. The sciatic nerve also harbors T cells of the gamma delta subtype which express *Trdc*, innate lymphoid cells (ILCs) and natural killer (NK) cells (Fig. 2b).

Particularly prominent was an increase in recruited macrophages (RecMacs) in nerves from mice on the HFHFD (Fig. 2c,2d,S4b,S6b). These cells had likely infiltrated from the bloodstream recently, as they expressed markers like *Ccr2*, *Fn1*, and *Ly6c2* but had acquired a macrophage identity expressing *Csf1r* and *Mrc1*. We used flow cytometry to independently validate the increase in recruited macrophages (Ly6C^hi^Cd64+Cd11b+ population) (Fig. 2e, S5, S6a). An increase in recMacs was first observed at 12 weeks in both male (Fig. 2e) and female mice (Fig S1d), which coincides with the onset of heat hypoalgesia, suggesting the macrophages may be recruited in response to either changes in neuronal activity or the nerve microenvironment. However, at this time on the diet (12 weeks), we found no evidence of terminal axonal loss in the skin (Fig 1h) or demyelination in the nerve (Fig S2c,S2d,S2e), suggesting that, unlike Wallerian degeneration, macrophage recruitment into diabetic nerves is not driven by axonal degeneration.

To examine if there are transcriptional changes in the resident sciatic nerve macrophages (resMacs) of diabetic mice, we conducted a differential gene expression analysis between these resMacs populations in HFHFD vs CD samples (*Csf1r+ Ccr2^-^* clusters) and found that resMacs from the HFHFD mice nerves upregulate many chemokines, including *Ccl3* and *Ccl8* (Fig. 2f). A gene ontology (GO) pathway analysis revealed that chemotaxis pathways are upregulated in the resident macrophages (Fig. S6c) suggesting they very likely recruit the recMacs population. We validated the increase in *Ccl2* mRNA in HFHFD nerves by RT-qPCR (Fig. 2g).

In the skin, we identified all expected immune cell types^23^, including macrophages, dendritic cells, Langerhans cells, T cells, mast cells, and ILCs (Fig. S7B). We evaluated the cell type proportions and found no evidence of any immune cell infiltration into the skin (Fig. S7c,S7d). This suggests that immune infiltration does not occur in the skin at this early time, indicating that macrophage infiltration into the nerve, is tissue specific rather than a systemic immune alteration, and is likely due to specific local changes in the nerves of diabetic mice.

### Macrophages recruited into diabetic nerves resemble those recruited after a sciatic nerve crush

To evaluate whether the macrophages recruited into the peripheral nerves of diabetic mice are similar to those that respond to axonal injury, we compared *Csf1r+* clusters obtained from nerves of mice on the HFHFD with a previously reported dataset from sorted sciatic nerve macrophages after a nerve crush injury^4^ (Fig. 3a). After merging the datasets, a clustering analysis revealed that both datasets contain similar cell types, including resident and recruited macrophage clusters. In addition, macrophages from the nerves of mice on the CD for 12 weeks and intact naive nerves showed a similar cellular composition, including mainly resident macrophage subtypes (Fig. 3b,d).

**Figure 3.**
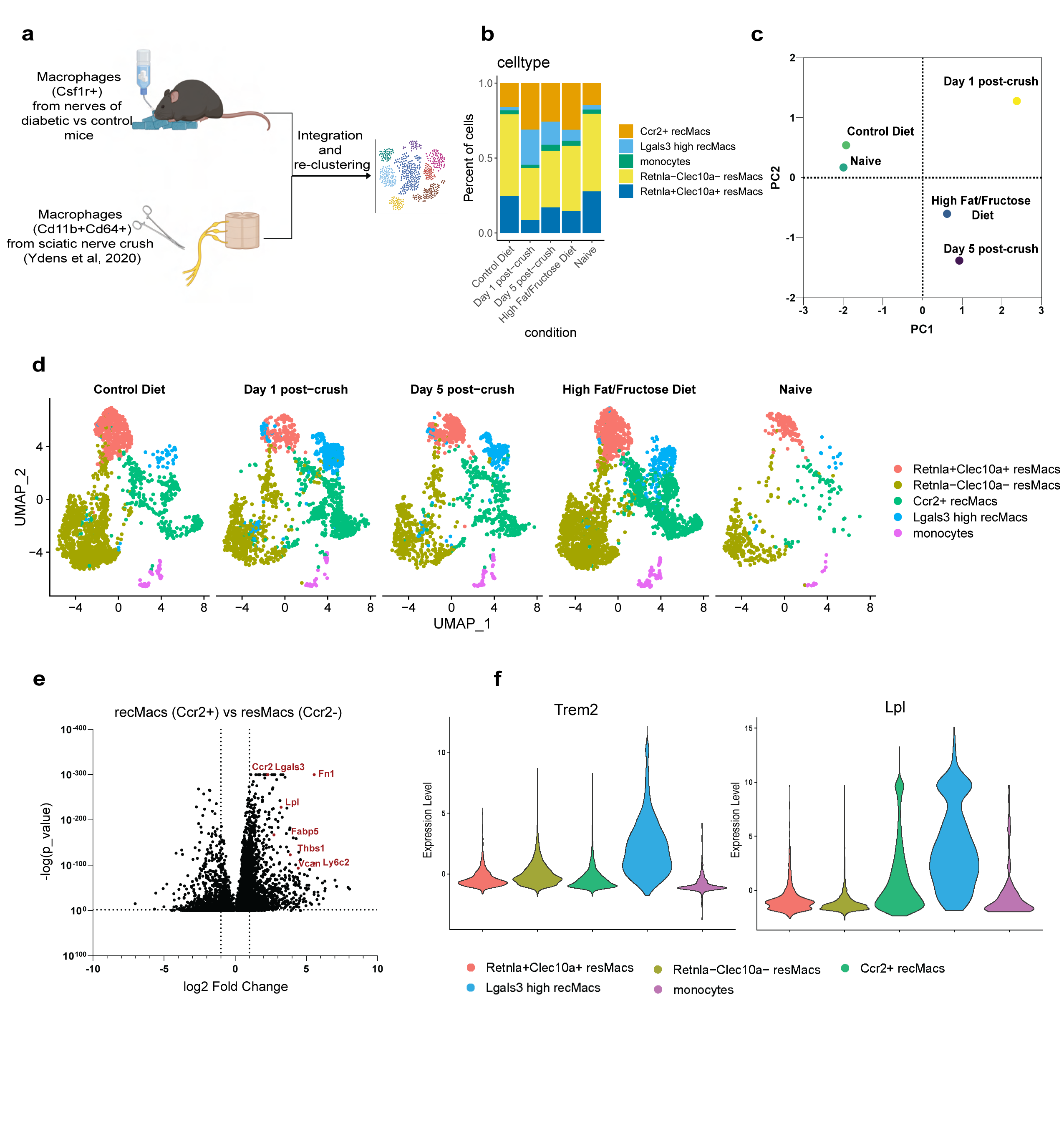
Macrophages recruited into nerves of diabetic mice after 12 weeks are similar to those recruited days after a sciatic nerve injury. **a.** Integration of macrophages from this dataset and a published nerve crush dataset. **b.** Cell proportions under the different conditions in the integrated datasets **c.** PCA plot of cell proportions. **d.** UMAP plots from merged dataset split by condition. **e.** Volcano plot of gene markers differentially expressed between CCR2+ macrophages (recMacs) and CCR2-macrophages (resMacs). **f.** Violin plots of an injury responsive gene (*Trem2*) and lipid-responsive gene (*Lpl*) in the macrophage datasets.

In terms of recruited macrophages, two clusters emerge in the integrated data, a cluster high in CCR2 (Ccr2+ recMacs) and a cluster high in *Lgals3* expression, (Lgals3 high recMacs) (Fig. 3d). Those two clusters were recruited in both the HFHFD and the post-crush (day 1 and day 5) samples (Fig. 3d). The recruited macrophages in both the HFHFD and nerve injury samples also expressed neurodegenerative disease associated microglial (DAMs)^5^ gene markers such as *Trem2* and *Lpl* (Fig. 3f). From the gene expression programs of the macrophages in diabetic nerves, we identified differentially expressed genes between recMacs and resMacs and found many injury-responsive genes such as *Fn1* and *Chil3*^4^, as well as genes responsible for lipid uptake such as *Lpl*^24–27^ and genes previously shown to promote nerve regeneration such as *Lgals3* and *Thbs1*^27–29^ (Fig. 4e). These data reveal that the macrophages that infiltrate the intact nerves of diabetic mice are similar to those macrophages recruited rapidly and distally after a nerve injury, even though the two insults are quite distinct, and express multiple similar genes that suggest a pro-repair phenotype.

**Figure 4.**
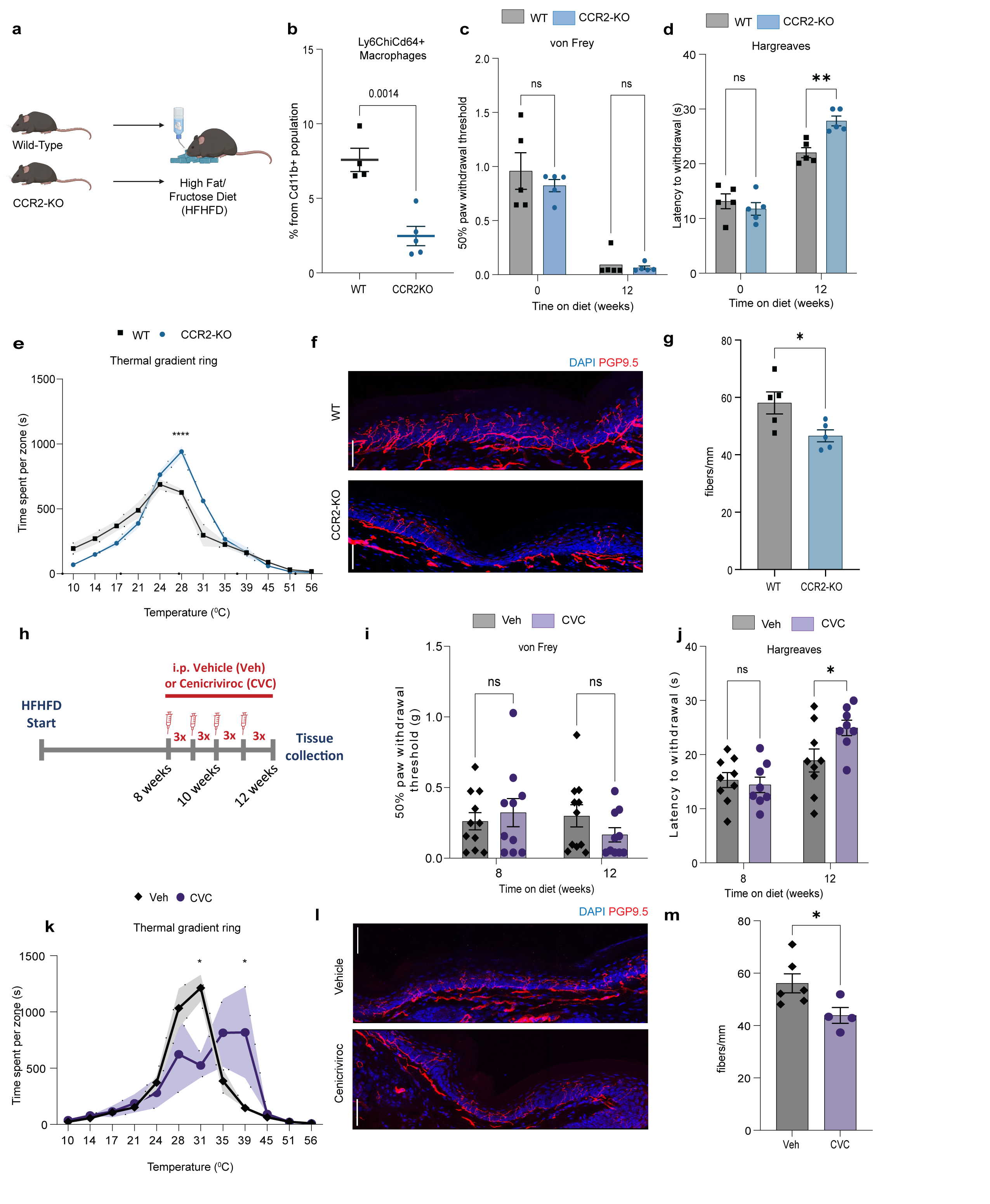
CCR2+ macrophages are neuroprotective and blocking their recruitment accelerates diabetic neuropathy. **a.** Illustration of experiment. **b.** Flow cytometry analysis of recruited macrophages (Ly6c+Cd64+Cd11b+) in sciatic nerves of CCR2-KO and WT mice fed the HFHFD for 12 weeks (p-value=0.0014). **c.** Mechanical withdrawal thresholds, **d.** Heat withdrawal latencies (p-value at 12w = 0.0037), **e.** Time spent per zone on thermal gradient ring (n=5 WT and n=4 KO mice) (p-value at 28w <0.0001), **f.** Representative PGP9.5 staining from hind paw skin **g.** Quantification of intraepidermal nerve fiber densities from CCR2-KO and WT mice fed a HFHFD for 12 weeks (p-value =0.0310). **h.** Timeline of CCR2/CCR5 inhibitor Cenicriviroc (CVC), injections starting after 8 weeks on diet. **i.** Mechanical withdrawal thresholds, **j.** Heat withdrawal latencies (p-value at 12w = 0.0303), **k.** Time spent in different temperature zones on thermal gradient ring(n=4 mice per group) (p-value at 31C = 0.0105, at 39C = 0.0135), **l.** Representative PGP9.5 staining from hind paw skin **m.** Quantification of intraepidermal nerve fiber densities from CVC and vehicle injected mice fed on HFHFD at 12 weeks (p-value = 0.0436).

### Recruited macrophages delay the onset of diabetic peripheral neuropathy

Recruited macrophages in injured peripheral nerves contribute to the clearance of both myelin and axonal debris, enabling axonal regeneration and recovery of function^30^. To evaluate whether macrophages recruited into the nerves of diabetic mice have a similar neuroprotective role, we fed CCR2-KO and WT mice on the HFHFD and evaluated whether preventing CCR2-dependent macrophage recruitment into the nerve had an impact on the axonal disease progression (Fig. 4a). The numbers of Ly6C^hi^ Cd64+ recruited macrophages in the sciatic nerve in CCR2-KO mice was reduced compared to WT mice, indicating that the macrophage recruitment in HFHFD mice is CCR2-dependent (Fig. 4b). CCR2-KO mice exhibited a greater heat hypoalgesia than age-matched WT mice (Fig. 4d), but no difference in mechanical allodynia (Fig. 4c). These mice also had more heat tolerance on the thermal gradient ring than their WT counterparts, confirming the diminished noxious heat sensitivity (Fig. 4e). CCR2-KO mice at 12 weeks on the HFHF diet also had a reduced IENFD, revealing a much earlier onset of terminal axonal neuropathy than in WT mice on the HFHFD (Fig 4f,4g). The CCR2-KO mice fed on the HFHFD for 12 weeks gained weight to the same extent as their WT counterparts (Fig. S8a).

To block CCR2 recruitment at a time after the onset of the disease we used the CCR2/CCR5 inhibitor Cenicriviroc (CVC)^31^. After assessing behavioral changes at 8 weeks, mice were randomly placed in CVC treatment or vehicle groups. Mice treated with CVC starting after 8 weeks on the diet (Fig. 4h) and which did not lose weight, were evaluated for behavioral and histological changes (Fig. S8b). Like the CCR2-KO mice, the CVC-injected mice had a persistent mechanical allodynia (Fig. 4i) and showed higher heat latencies compared to the vehicle treated group at 12 weeks of diet (Fig. 4j). They also had a greater heat tolerance on the thermal gradient test (Fig. 4k) and detectable axonal loss in the epidermis at 12 weeks, consistent with a more severe and earlier onset peripheral axonal disease, as with the CCR2-KO mice (Fig. 4l,4m). These data reveal that a CCR2 dependent recruitment of macrophages into diabetic peripheral nerves both reduces the severity of behavioral sensory loss and delays the progression of axonal degeneration in the skin. In consequence, we conclude that blocking the recruitment of macrophages into peripheral nerves in type 2 diabetes is detrimental and that promoting and sustaining this recruitment is beneficial.

## Discussion

We have utilized a diet-induced model of type 2 diabetes and found that macrophages with an expression profile resembling neurodegenerative disease-associated myeloid cells, infiltrate the intact peripheral nerves of diabetic mice, and that their recruitment is neuroprotective and delays the onset of terminal axonal degeneration. About 50% of patients with diabetes develop diabetic neuropathy, half of which experience pain^3^. In the HFHFD model, mechanical allodynia developed early in the disease (3 months) but then resolved by six months. A transient mechanical allodynia is also observed in a genetic model of leptin deficiency (db/db)^32^. Preventing axonal degeneration by Sarm1-KO was associated with a persistence of the allodynia, suggesting that blocking degeneration does not eliminate pain hypersensitivity. Heat hypoalgesia manifested in HFHFD treated mice well before axon terminal degeneration was detectable, but preventing axon terminal degeneration by Sarm1 knockout prevented this lack of sensitivity to noxious heat. Patients often present with sensory abnormalities years before a definitive skin biopsy diagnosis of axonal loss suggesting the mouse model captures major features of the human disease^14,33^. What is responsible for the early, pre-neuropathy heat hypoalgesia now needs to be determined and why it has a SARM1 dependent component in the absence of detectable axon degeneration.

We find that macrophages are recruited into peripheral nerves but not the skin of diabetic mice. Mice on a HFD present with an upregulation of *Lgals3* in the sciatic nerve^22^, which is one of the most highly expressed genes in the recMacs that we find infiltrate into the nerves of HFHFD diabetic mice (Fig.4f). *Lgals3* encodes Galectin-3, whose knockdown delays axonal regeneration after nerve injury^28,34^ and is a Trem2 ligand in microglia reacting to neurodegeneration^35^. Galectin-3-expressing phagocytic macrophages infiltrate into diabetic brains^36^ suggesting that the peripheral nerve infiltrating macrophages may also be phagocytic, and that they may be neuroprotective by virtue of clearing debris or excess fatty acids in the nerve. Galectin-3 is expressed by microglia at sites of brain atrophy and is protective against retinal degeneration^29^. Another gene highly expressed by the nerve infiltrating macrophages encodes Thrombospondin-1, another neuroprotective mediator^27^. Thrombospondin-1 is also an inhibitor of the peripheral sensitization of nociceptors^23^. Similarities observed between the recMacs in diabetic peripheral nerves and the recMacs after nerve injury as well as disease-associated microglia (DAMs)^5^ indicate that there is a common myeloid cell identity in quite different neurological diseases which may reflect similar neuroprotective roles.

Macrophages may be recruited into peripheral nerves early in diabetes in response to changes in the nerve microenvironment. Non-neuronal cell types, such as Schwann cells could be modified by the HFHFD, as they are responsive to changes in glucose or insulin^22^ and may release mediators that alter nerve resident macrophages such that they recruit circulating myeloid cells. Metabolomic studies show that a HFD alone and when supplemented with low dose streptozotocin modifies the fatty acid composition of the nerve^37^. Perhaps at 12 weeks on a HFHFD an excess of free fatty acids exerts effects on nerve resident macrophages similar to that exerted by membrane or myelin debris in acutely degenerating nerves, which would explain the similar recruited macrophage response after a nerve crush and in diabetes. A GO pathway analysis of the resident macrophages in the nerves from our diabetic mice, revealed “fatty acid transport” as one of the enriched pathways (Fig. S6c), a finding compatible with a response to an excess of fatty acids. Increased fatty acid oxidation and uptake can promote macrophage differentiation into reparative identities^38^.

Although macrophages have long been considered to have a pathogenic role in diabetes and insulin resistance^39,40^, we now find that early recruited macrophages in nerves delay the onset of neuropathy and reduce its extent. While macrophages were suggested to play a detrimental role in a HFD-induced model of neuropathy^41,42^, these studies only focused on mechanical allodynia and did not investigate either heat sensitivity or skin innervation. We find that mechanical allodynia is transient in our mouse model of type 2 diabetes, and its resolution may be driven by the degeneration of mechanosensitive fibers rather than an improvement in disease progression, as supported by our results in Sarm1-KO mice. In addition, mechanical allodynia is the least consistent change associated in diabetic patients whereas histological changes in the skin revealing axon terminal loss are the gold standard for the diagnosis of DPN^3,14^ Therefore it is important that mouse models also exhibit this feature of the disease. Because the transcriptional profile of macrophages recruited to diabetic nerves are similar to those found in nerve crush-induced macrophages, and because their role is, we find, neuroprotective, it will be important to determine whether a similar protection occurs in non-diabetic peripheral neuropathies. It also will be worthwhile to determine whether macrophage recruitment in peripheral nerves correlates with a better prognosis in diabetic patients for not developing neuropathy. Our findings provide a novel insight into the neuroprotective role of immune cells in diabetic neuropathy that can now drive exploration of biomarkers to predict disease progression as well as develop therapeutic strategies that maximize and sustain the neuroprotective action of macrophages recruited into nerves.

## Methods

### Mice

8-12 week old C57Bl/6J mice were obtained from the Jackson Laboratory (JAX: 000664). CCR2-KO (strain no:004999) and Sarm1-KO (strain no:018069) mice were obtained from Jackson Laboratory. Both male and female mice were used unless otherwise specified. Single-cell RNA sequencing was performed on male mice. All animal experiments and procedures were conducted according to the institutional animal care and safety guidelines at Boston Children’s Hospital.

### Diet

High Fat Diet containing 60% Fat from Lard was obtained from research diets (**D12492**) along with a matched sucrose Control Diet (**D12450J**). Diet experiments were started at 10-12 weeks of age. Mice that were given the HFHFD were also given water bottles with 42g/L D-(-)-fructose in autoclaved water and those given CD were given plain autoclaved water. Mice food and water levels were monitored 3x per week and replenished as needed throughout the entirety of their diet regimen.

### Glucose Tolerance test

Mice were fasted overnight, and baseline blood glucose was measured from a tail nick using One Touch Ultra 2 glucose monitoring system. Mice were then injected with 5uL/g weight of 0.2g/mL glucose i.p. and blood glucose was measured at 15, 30, 60, and 120 minutes.

### Behavior

#### Von Frey

Mice were habituated to von Frey chambers and mesh bottom for 2 days, 1 hour each day, prior to testing. On the day of the test, mice were habituated for 1h before the test. The up-down method was employed to determine the mice’s 50% withdrawal threshold^43^. Behavioral experiments to compare CD and HFHFD were difficult to conduct in a blinded fashion since mice weights and feces colors were obvious; however, these were conducted by at least 2 independent investigators to validate the findings. All subsequent experiments with genetic or pharmacological manipulations were conducted in a blinded fashion (this was also done for the Hargreaves test) ***Hargreaves***. Mice were habituated to the Hargreaves apparatus (IITC #390G) consisting of a glass floor heated to 30^0^C and a plexiglass chamber for 2 days prior to testing, 30 minutes each day. On the day of the assessment, mice were habituated for 1 hour before the test. A focused heat light source was shined on the plantar surface of the left paw of the mice and a ramping heat stimulus was therefore applied until paw withdrawal was recorded. Readings were averaged from 3 trials per mouse.

#### Thermal Gradient Ring

Mice were placed in the thermal gradient ring (Ugo Basile, 35550) which was set to 10^0^C on one side and 56^0^C on the other side. Their location, movement, and temperature of the plates were recorded for an hour. All mice in a single experiment were recorded using one apparatus. Using the ANY MAZE software, the time spent per zone as well as distance traveled were recorded for each mouse.

### Skin immunohistochemistry

A 2mm punch from the hind paw plantar skin of mice was collected in Zamboni Fixative (Newcomer supply, #1459A) overnight in 4C on a piece of filter paper to flatten the tissue. After fixation, skin samples were kept in 30% sucrose for at least 48 hours in 4C. Skin samples were then frozen in OCT, cryosectioned at 30um thickness and mounted onto superfrost plus microscope slides. For immunostaining, slides were thawed for 1h at RT then washed in PBS for 15 minutes following 3 10-minute 1% Triton-X washes. Slides were incubated in blocking buffer for 2h at RT (10% Donkey Serum, 0.4% Triton-X, 0.05% Tween 20, 1% BSA) and then incubated in primary antibody overnight at 4C (Rabbit anti-PGP9.5 Abcam cat# 1:200). The next day, slides were washed 3x with PBS and incubated in secondary antibody for 2h at RT (Donkey anti-Rabbit Cy3 Jackson ImmunoResearch 711-165-152 1:500). Slides were then washed 3x with PBS and mounted with Prolong antifade DAPI medium (Invitrogen, #P36935). Confocal images were taken on a Leica SP8 microscope with a 63x oil objective. A z-stack spanning the 30um thickness was taken and 4 adjacent images were tiled. Using ImageJ, maximum intensity projection was taken, and the number of fibers reaching the epidermis was counted and divided by the length of the epidermal layer in the image to obtain an IENFD value for that image. At least 2 images of non-consecutive sections were averaged per mouse to obtain the IENFD value for that mouse.

### Sciatic nerve tissue collection and processing

Sciatic nerves were dissected and collected in ice cold 1%BSA/RPMI. Tissues were moved for digestion in Collagenase A (Roche, 5mg/ml) and DispaseII (Roche, 1mg/ml) and minced before placing on thermocycler at 37C for 1h at 1000rpm. Samples were then filtered using a 70um filter, washed in FACS buffer (2% FBS, 2mM EDTA, PBS) then washed with 1x PBS to obtain single cell suspensions.

### Flow Cytometry

Cells were then incubated for 10min at 4C in Fc Block (Tonbo Biosciences cat # 70-0161-U500, 1:2000) then washed with FACS buffer. Samples were then stained for 30 minutes with antibodies against mouse CD45-ef780 (Thermo Fisher scientific, clone 30-11, cat #47-0451-82, 1:400), CD11b-FITC (Thermo Fisher Scientific, clone M1/70, cat #11-0112-82, 1:400), CD64-PE-594 (BioLegend, clone X54-5/7.1, cat #139319, 1:400), Ly6c-BV711 (BioLegend, clone HK1.4, cat #128037,1:1000), Ly6g-PE (BioLegend, clone 1A8, cat #127607, 1:800), for 30 min on ice then washed with FACS buffer twice and resuspended with 3uM DAPI before analyzing on a LSR Fortessa Cytometer and analyzing using FlowJo.

### Preparation of sciatic nerve immune cells for single cell sequencing

Single cell suspensions collected as described above from 2 mice (4 sciatic nerves) were pooled in each sample. Samples were resuspended in FACS buffer (2% FBS, 2mM EDTA, PBS) and then incubated for 10min at 4C in Fc Block (Tonbo Biosciences cat # 70-0161-U500, 1:2000) and washed with FACS buffer. Cells were then stained for 30min in anti CD45 conjugated APC-Cy7 antibody (Thermo Fisher scientific, clone 30-11, cat #47-0451-82, 1:200) in FACS buffer then washed 2x in 0.5%BSA/PBS solution with no EDTA. Cells were then resuspended with 3uM DAPI in the 0.5%BSA/PBS solution before Live Cd45+ cells were sorted on a BD FACS Aria sorter and processed for 10x sequencing following 10x protocol.

### Barcoding and library preparation

The Chromium Next GEM Single Cell 3′ Reagent kit v3.1 (Dual Index) was used. Barcoding and library preparation was performed following the manufacturer’s protocols. Briefly, to generate single-cell gel-bead-in-emulsion (GEMs) solution, sorted cells were resuspended in a final volume of 40 μl and were loaded on a Next GEM Chip G (10X Genomics) and processed with the 10X Genomics Chromium Controller. Reverse transcription was performed as instructed: 53°C for 45 min and 85°C for 5 min in a thermocycler. Next, first-strand cDNA was cleaned with DynaBeads MyOne SILANE (10 X Genomics, 2000048). The amplified cDNA, intermedium products, and final libraries were prepared and cleaned with SPRIselect Regent kit (Beckman Coulter, B23318). A small aliquot of each library was used for quality control to determine fragment size distribution and DNA concentration using a bioanalyzer. Libraries were pooled for sequencing with a NovaSeq 6000 (Illumina) at an estimated depth of 110,000 reads per cell. Sequencing reads were aligned to the mouse reference transcriptome (mm10, version 2020-A) using CellRanger (v.7.0.1).

### scRNAseq data analysis

Raw scRNAseq data were processed using the 10X Genomics CellRanger software version 7.0.01. The CellRanger ‘mkfastq’ function was used for de-multiplexing and generating FASTQ files from raw BCL. The CellRanger ‘count’ function with default settings was used with the mm10 reference supplied by 10X Genomics, to align reads and generate single-cell feature counts. Ambient RNA contamination was removed from each sample using CellBender6 (‘remove-background’, default parameters). Seurat version 5.0.1 implemented in R version 4.3.2 was used for downstream analysis. Cells were excluded if they had fewer than 500 features, or fewer than 250 genes, or the mitochondrial content was more than 20%. 2 CD and 2 HFHFD samples were integrated and normalized following previously described Seurat SCTransform+ CCA integration pipeline^44^. The mitochondrial mapping percentage was regressed out during the SCTransform normalization step. PCA was performed and the top 40 principal components were used for downstream analysis. A K-nearest-neighbor graph was produced using Euclidean distances. The Louvain algorithm was used with resolution set to 0.4 to group cells together. Nonlinear dimensional reduction was done using UMAP. FindAllMarkers function was used to determine marker genes for each cluster and clusters were identified based on canonical markers previously published in the literature^4,11,23^. Differentially expressed genes were determined using the FindMarkers function which performs differential expression testing based on the non-parametric Wilcoxon rank sum test.

### Integration with macrophages after nerve crush dataset

All clusters expressing *Csf1r* were subsetted into new dataset and raw data file from Ydens et al^4^ was downloaded. Samples from 2 CD, 2 HFHFD were merged using Seurat with Ydens’ Naïve, Day 1 post-crush and day 5 post-crush samples. Data was filtered according to the same filters above and integrated using SCTransform as described above. Dataset was then re-clustered with 0.4 resolution and FindAllMarkers was used to identify cell types.

### Nerve electron microscopy

Sciatic nerves were collected in 2.5%Glutaraldehyde,2.5%Paraformaldehide in 0.1M sodium cacodylate buffer (pH7.5) and fixed in the same solution overnight. Small pieces (1-2mm) of fixed tissue were washed in 0.1M cacodylate buffer and postfixed with 1% Osmiumtetroxide (OsO4)/1.5% Potassium ferrocyanide(KFeCN6) for 1 hour, washed in water 2x, then washed 1x in 50mM Maleate buffer pH 5.15 (MB) and incubated in 1% uranyl acetate in MB for 1hr followed by 1 wash in MB, 2 washes in water and subsequent dehydration in grades of alcohol (10min each; 50%, 70%, 90%, 2x10min 100%). The samples were then put in propyleneoxide for 1 hr and infiltrated ON in a 1:1 mixture of propyleneoxide and TAAB Epon (TAAB Laboratories Equipment Ltd). The next day, samples were embedded in TAAB Epon and polymerized at 60 degrees C for 48 hrs. Ultrathin sections were cut on a Reichert Ultracut-S microtome, picked up on to copper grids, stained with lead citrate and examined in a JEOL 1200EX Transmission electron microscope or a TecnaiG² Spirit BioTWIN. Images were recorded with an AMT 2k CCD camera. Electron Microscopy Imaging, consultation and services were performed in the HMS Electron Microscopy Facility. G-ratios and % myelinated axons were calculated from the EM images using MyelTracer as described before^45^.

### Mouse injections

The CCR2/CCR5 inhibitor Cenicriviroc (Selleckchem, cat#S8512) or DMSO were diluted in corn oil and injected i.p. at 20mg/kg 3x per week starting 8 weeks on the HFHFD till 12 weeks. Mouse weights were recorded weekly, and mice who lost weight as a result of the drug treatment were excluded from the study before proceeding with behavioral testing because this counteracted the weight gain expected from HFHFD feeding.

### RT-qPCR

Mouse sciatic nerves were collected in RNAlater and stored at 4C overnight before long-term storage in −80C. To isolate RNA, nerves were moved from RNAlater to RLT buffer (Qiagen) and minced. Handheld homogenizer was then used to homogenize the nerves in RLT buffer and equal volume 70% ethanol was then added to the tube. RNA was isolated following Qiagen mini kit protocol. cDNA was synthesized from resulting RNA using cDNA VILO kit (Invitrogen, Cat#11754050). Relative gene expression was determined using gene-specific primers (PrimerBank) and SYBR Green master mix (Life Technologies) on a 7500 Fast Real-time PCR system (Applied Biosystems). Expression levels were normalized to *Hprt* levels using the 2–ΔCt method. The following primer sequences (5′-to-3′) were used: *Hprt* forward: CAGTCCCAGCGTCGTGATTA, *Hprt* reverse: TGGCCTCCCATCTCCTTCAT; *Ccl2* forward: GATGCAGTTAACGCCCCACT, *Ccl2* reverse: GAGCTTGGTGACAAAAACTACAGC.

### Statistical analysis

Statistical analysis was performed using GraphPad Prism Software. Two-group comparisons were made using two-tailed unpaired Student’s t-test. For comparisons of 2 groups over time, two-way ANOVA with appropriate multiple comparisons tests was used and 0.05 was set as the threshold for significance. Plots from behavior assays, immunohistochemistry and flow cytometry are representative of at least two independent repetitions.

## Acknowledgements

We thank Daniel Taub, Nick Andrews, Masakazu Kotoda, and Jacob Hickey from the Woolf lab for assistance. We thank Stephen Liberles, Maria Lehtinen, and Ruaidhri Jackson for thoughtful discussions. We thank Natalia Biscola and Leif Havton for the IHC protocol. We thank Mark Scimone and the Boston Children’s Hospital (BCH) Cellular Imaging Core; Ronald Mathieu and the BCH Flow Core; Maria Ericsson and the HMS Electron Microscopy Core; and the Technology Center for genomics and bioinformatics (TCGB) at UCLA. This work was supported by the National Institutes of Health (grant numbers F31NS127357 to S.H., R35NS105076 to C.J.W.), and the Dr. Miriam and Sheldon G. Adelson Medical Research Foundation (C.J.W and R.K.).

## Author Contribution

S.H. and C.J.W. conceptualized, led the design and execution of the study. S.H. performed behavioral assays with help from A.J., V.P., J.I. and E.S.D. A.J. and V.P. performed mouse injections. S.H. performed flow cytometry. SSA performed GTTs. S.H. performed single cell sequencing and analysis with help from A.J., J.I., R.K., Q.W., and D.H.G. S.H., J.I., E.S.D., D.N. performed IHC and IENFD analysis. S.H., J.I., E.S.D., D.N., and C.A.G. performed feeding experiments and weight monitoring. S.H., A.J., and C.J.W. guided experimental design and analysis. S.H. and C.J.W. wrote manuscript with input from all authors.

## Data availability

The raw data that support the findings are available from the corresponding author upon reasonable request.

## Supplementary Figure legends

**Fig S1.**
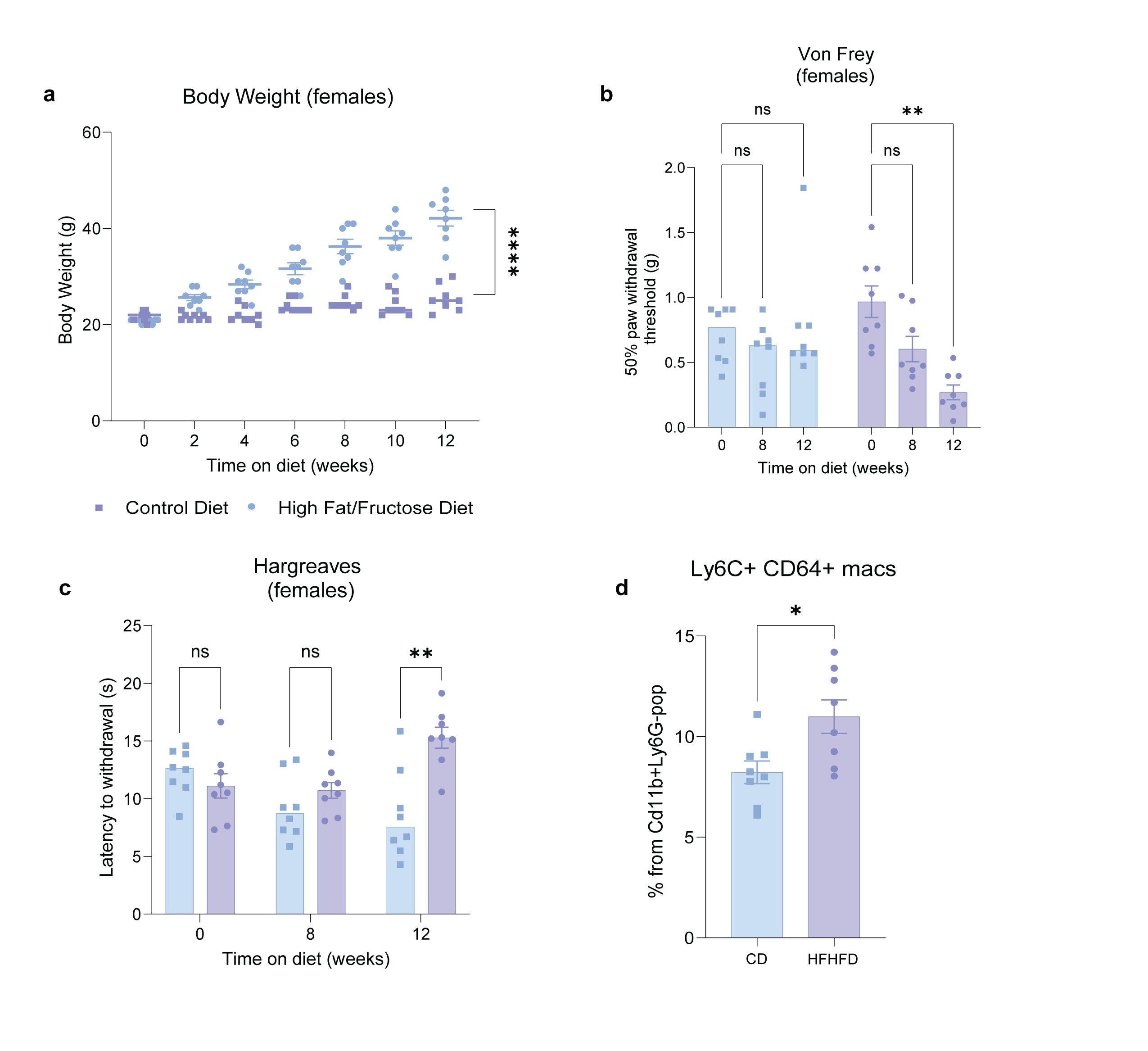
Female mice on HFHFD are equally susceptible to weight gain, behavioral changes, and immune cell infiltration in nerves. **a.** Body weight measurement over time on HFHFD vs CD feeding from female C57Bl6 mice (p-value <0.0001). **b.** Von Frey up down method was used to assess mechanical sensitivity in female mice fed HFHFD vs CD over 12 weeks (p-value 0 vs 12w = 0.0058). **c.** Hargreaves assay was used to assess heat sensitivity in female mice fed HFHFD vs CD over 12 weeks (p-value = 0.0044). **d.** Proportions of Ly6c+ CD64+ recruited macrophages from Live Cd45+ Cd11b+ cells in sciatic nerves of female mice fed HFHFD vs CD for 12 weeks (p-value = 0.0153).

**Fig S2.**
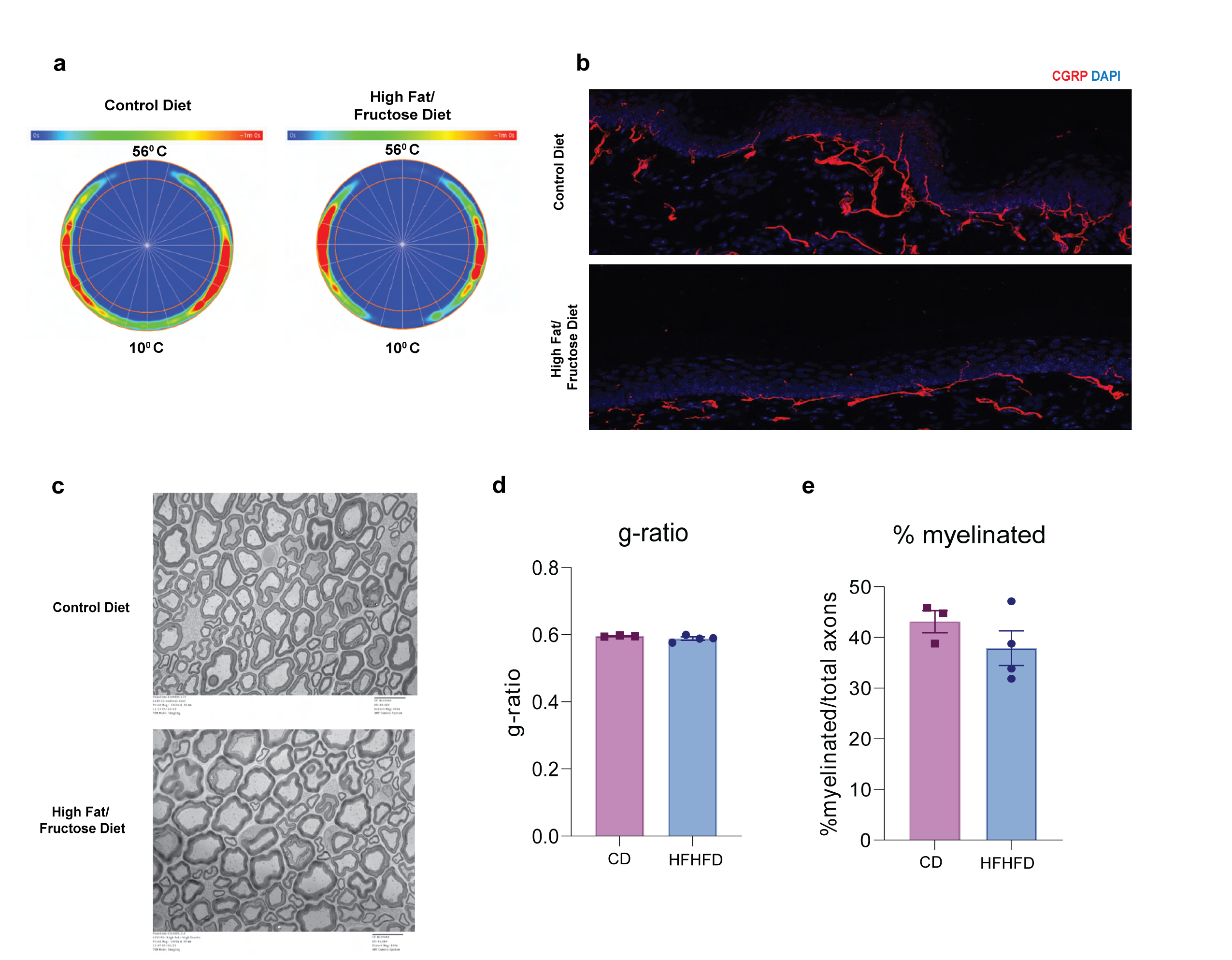
Behavior and histological parameters tested on CD vs HFHFD fed mice. **a.** Representative heat maps showing the location of the mouse during a 1-hour thermal gradient ring recording. **b.** Representative immunohistochemistry images with CGRP stained in red and DAPI in blue from CD and HFHFD hind paw skin after 24 weeks of feeding. **c.** Representative EM images from sciatic nerves of mice fed CD and HFHFD for 12 weeks. **d.** Quantification of EM images of g-ratio and **e.** percent myelinated fibers. Each dot represents an average of 2 images from each mouse.

**Fig S3.**
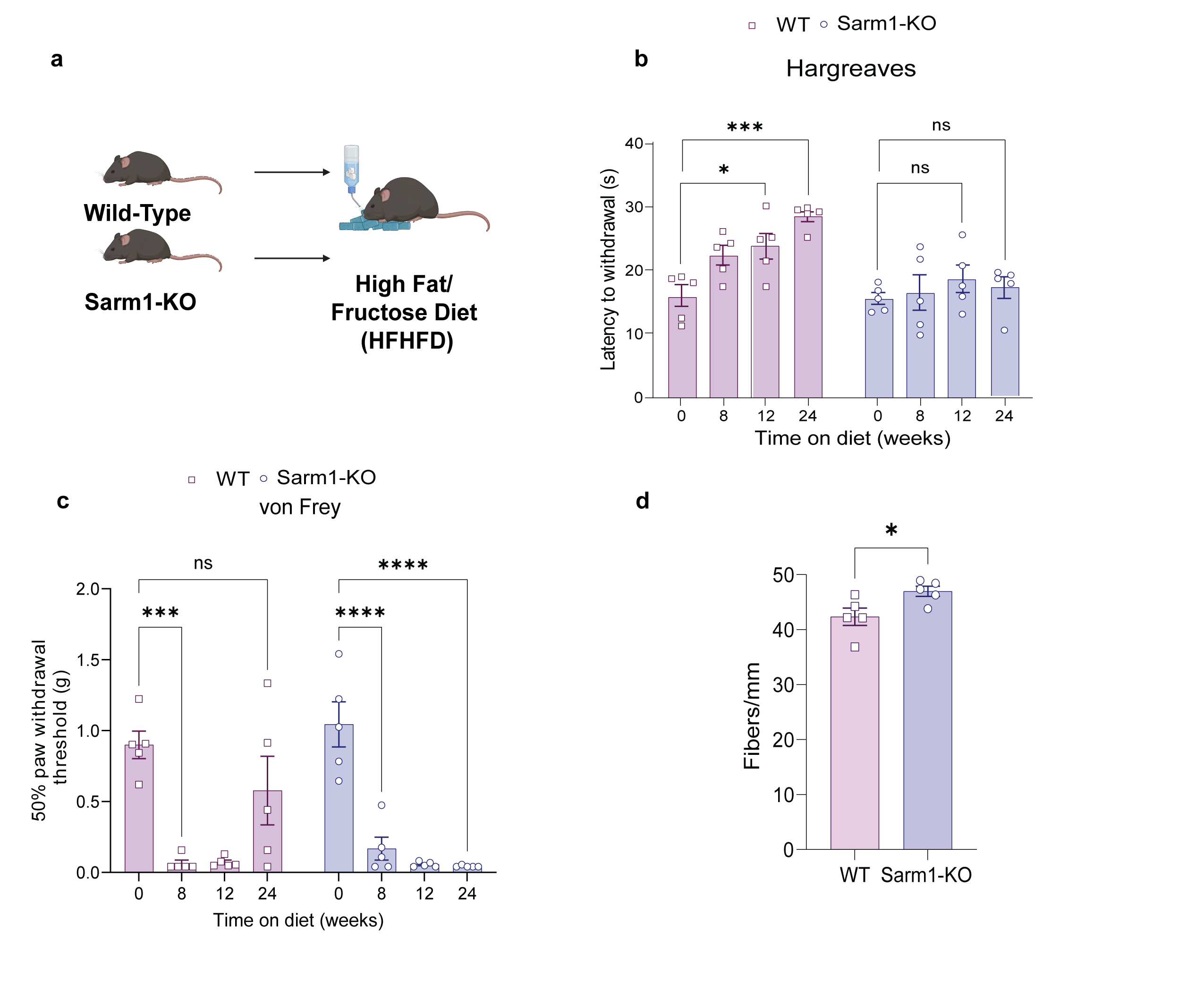
Sarm1 knockdown prevents the loss of heat sensitivity and skin axonal degeneration in HFHFD-fed mice but leads to persistent mechanical allodynia. **a.** Illustration of the experiment. **b.** Hargreaves test was used to assess thermal sensitivity in WT vs Sarm1-KO mice fed HFHFD or CD over 24 weeks. **c.** Von Frey filaments were used to assess mechanical sensitivity in WT vs Sarm1-KO mice fed HFHFD or CD over 24 weeks. **d.** Quantification of IENFD from PGP9.5 staining in hind paws of skin from WT and Sarm1-KO mice fed HFHFD or CD for 24 weeks (p-value = 0.0353).

**Fig S4.**
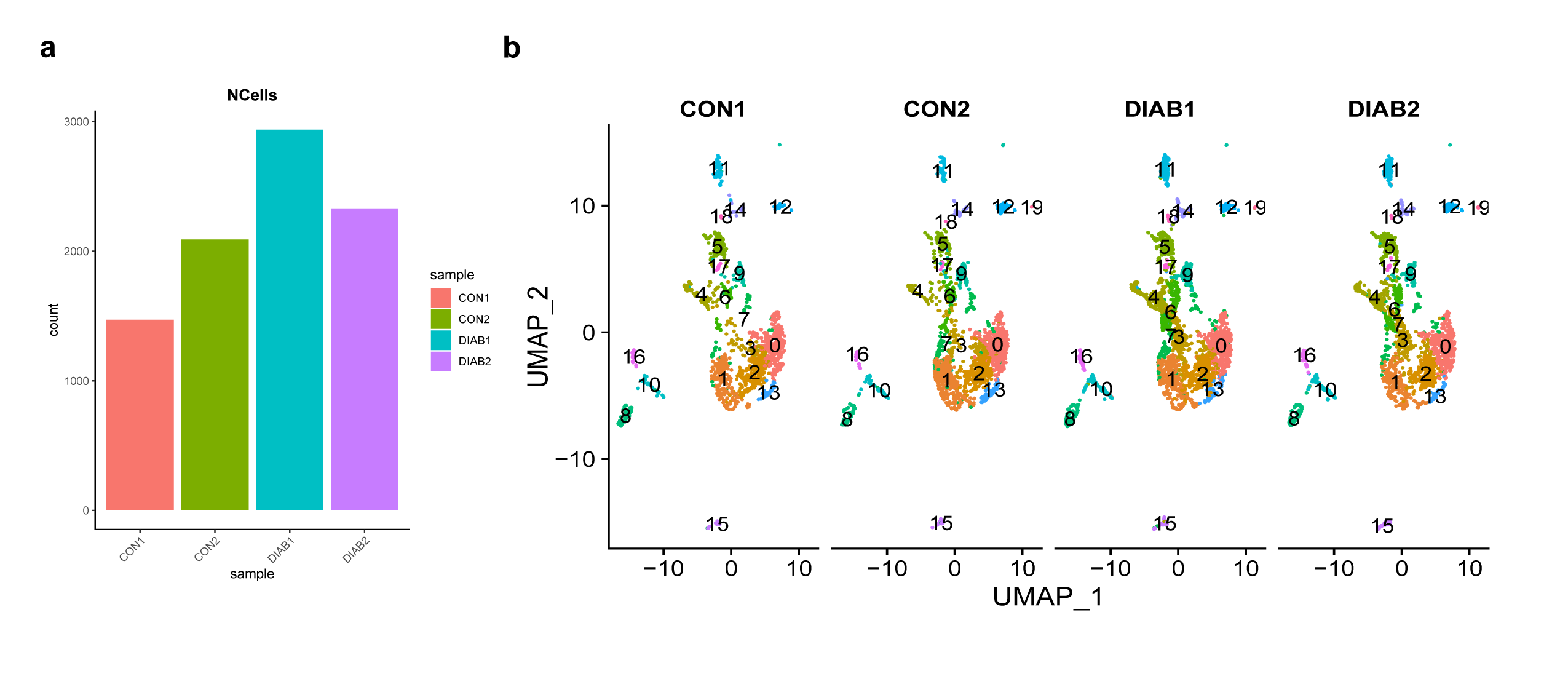
Sciatic nerve immune cell single cell sequencing. **a.** Number of cells analyzed in each sample post-filtering. **b.** UMAP plot showing similar cell composition in replicates of each condition.

**Fig S5.**
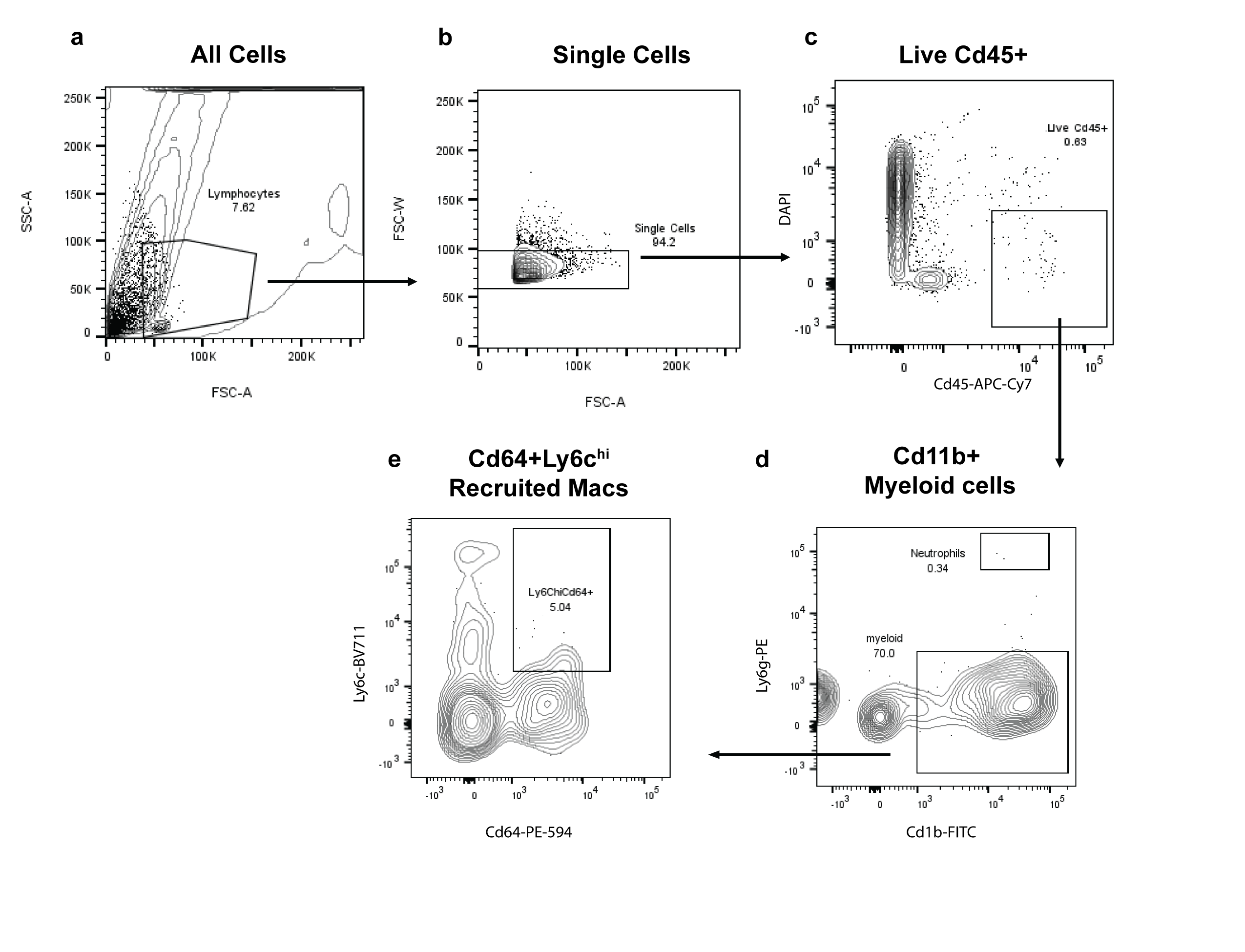
Gating strategy for flow cytometry assessment of recruited macrophages into the sciatic nerve. **a.** FSC-A and SSC-A were used to gate on all cells and exclude aggregates and small debris. **b.** FSC-A and FSC-W were used to gate on single cells. **c.** DAPI and CD45-APC-Cy7 were used to capture Live CD45+ immune cells. **d.** Cd11b-FITC and Ly6g-PE were used to gate on myeloid cells and exclude neutrophils. **e.** Cd64-PE-594 and Ly6c-BV711 were used to identify Ly6C+ Cd64+ recruited macrophages.

**Fig S6.**
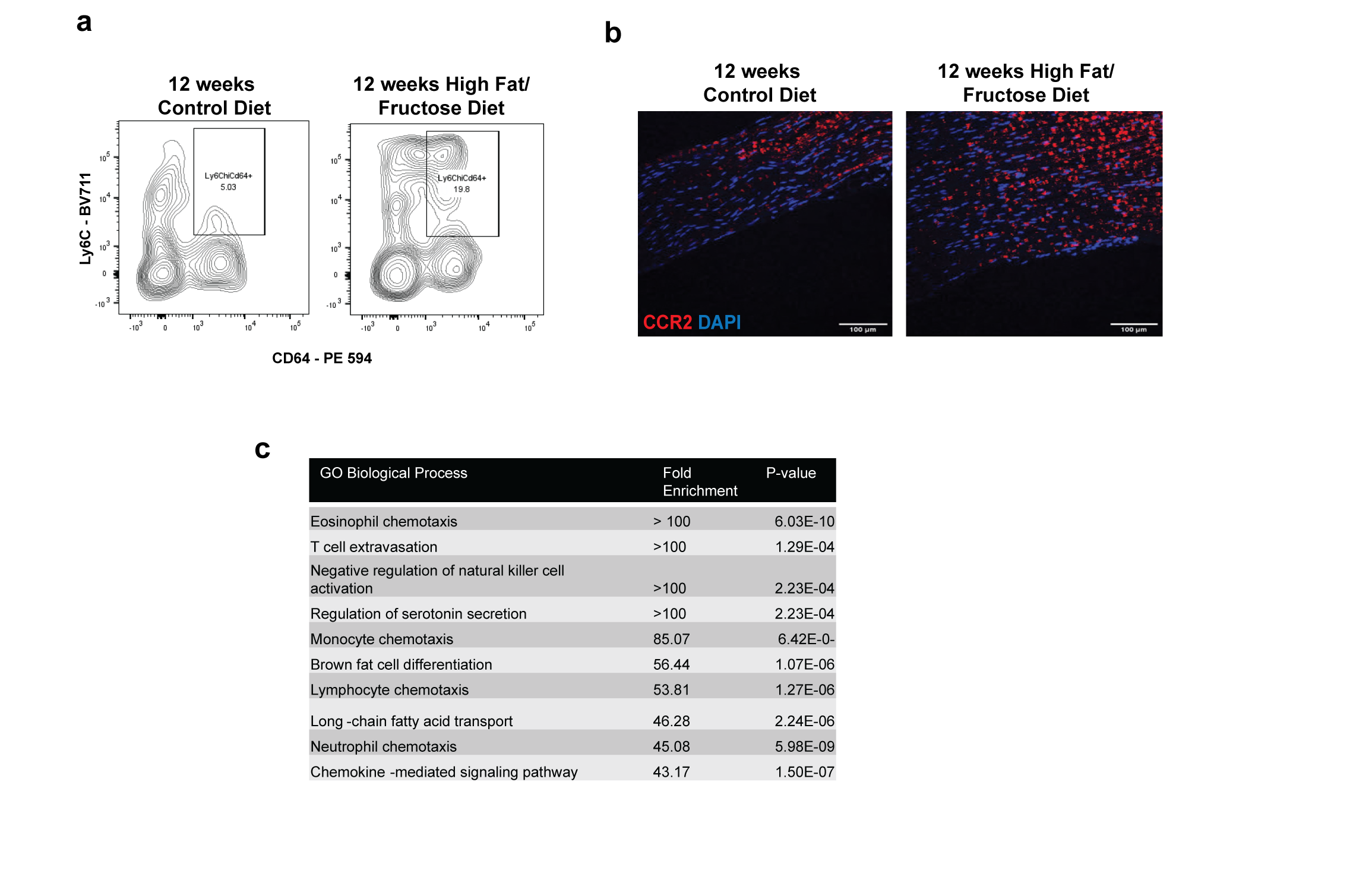
Changes in recruited and resident macrophages in sciatic nerves of mice fed HFHFD vs CD for 12 weeks. **a.** Representative flow cytometry contour plot of Ly6C and CD64 gate, pre-gated on Live CD45+Cd11b+ showing difference between CD and HFHFD sciatic nerves. **b.** Immunohistochemistry of sciatic nerves showing CCR2 in red, labeling recruited macrophages, and DAPI in blue from mice fed HFHFD vs CD for 12 weeks. **c.** GO pathway analysis from differentially expressed genes in resident resMacs clusters in HFHFD vs CD showing enrichment in chemotaxis pathways.

**Fig S7.**
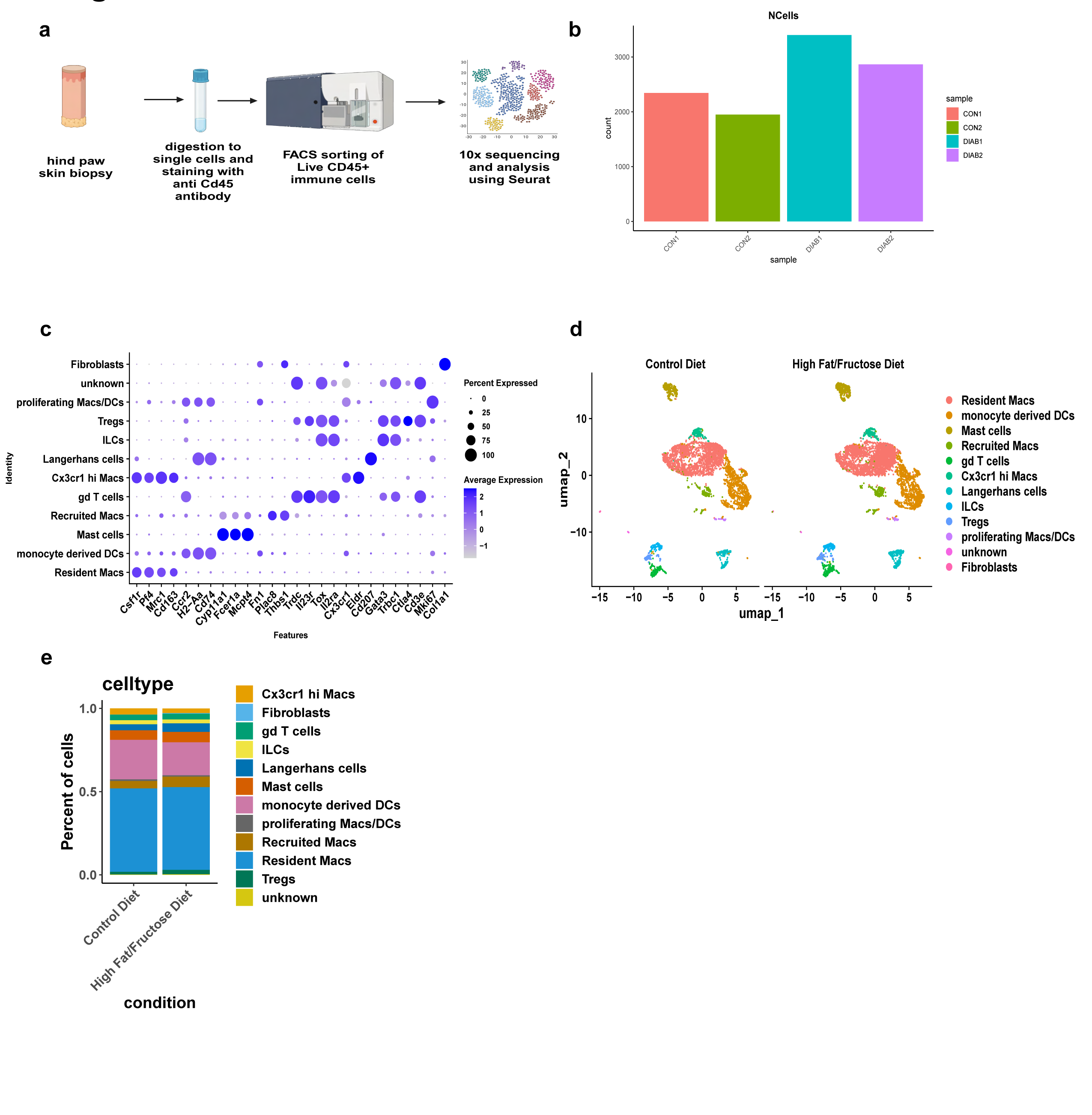
Single cell sequencing of skin immune cells from mice fed HFHFD or CD for 12 weeks shows no change in immune cell composition. **a.** Workflow for tissue collection, processing, sorting, and sequencing of skin immune cells from mice fed HFHFD or CD for 12 weeks from 4 mice per group pooled into 2 samples each group. **b.** Number of cells analyzed in each sample post-filtering. **c.** Dot plot of marker genes used to identify different immune cells in dataset. **d.** UMAP plot showing clusters identified in sciatic nerves of CD and HFHFD fed mice from 2 samples per group. **e.** Barplot showing proportions of the different cell types per group.

**Fig S8.**
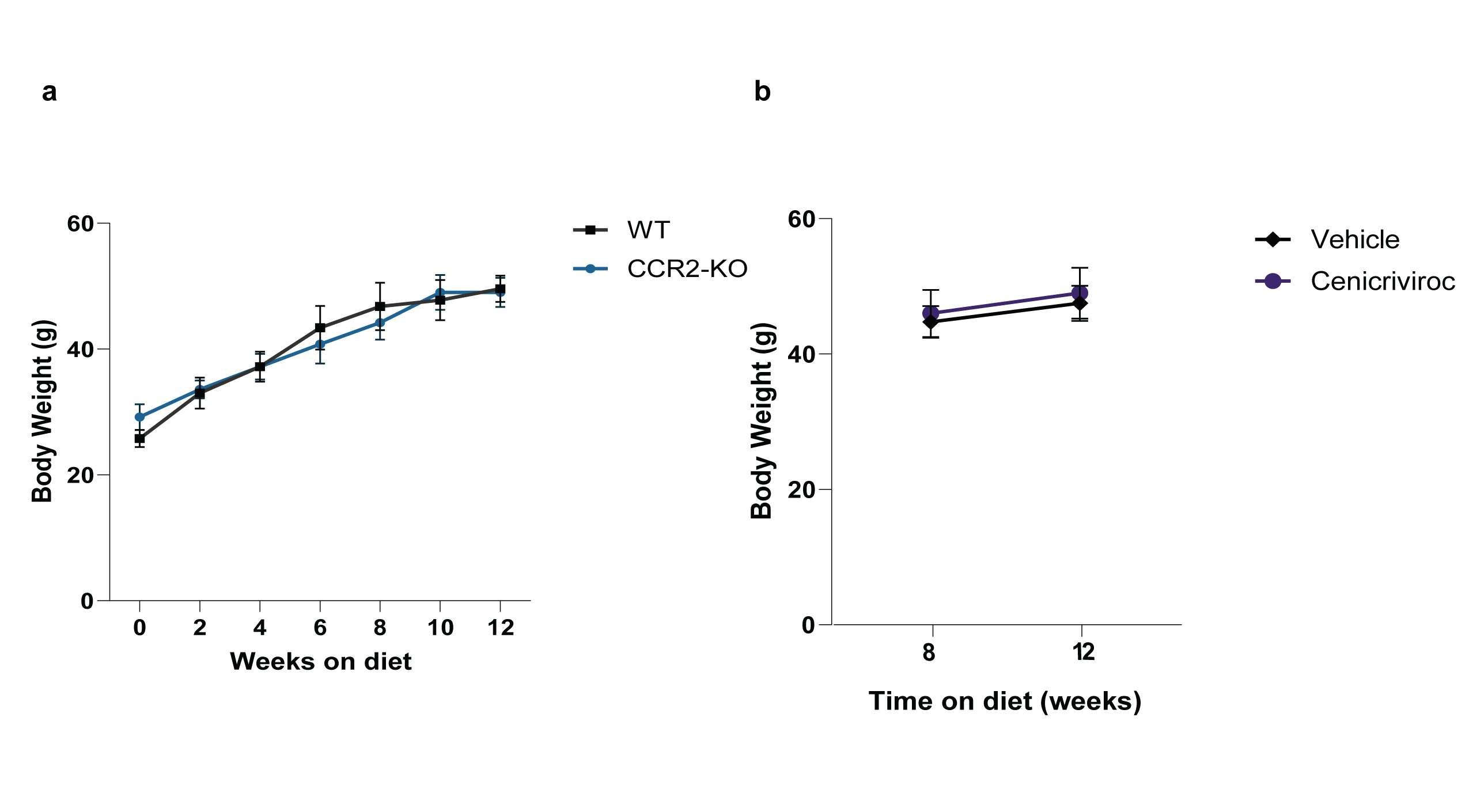
Blocking macrophage recruitment does not affect body weight gain. **a.** Body weight measurements in age matched CCR2-KO and WT mice over 12 weeks on HFHFD (n=5 mice per group). **b.** Body weight measurement of mice injected with CVC or vehicle before start of injections and at the end of experiment (n=4 per group).

